# Prior expectations induce pre-stimulus sensory templates

**DOI:** 10.1101/119073

**Authors:** Peter Kok, Pim Mostert, Floris P. de Lange

## Abstract

Perception can be described as a process of inference, integrating bottom-up sensory inputs and top-down expectations. However, it is unclear how this process is neurally implemented. It has been proposed that expectations lead to pre-stimulus baseline increases in sensory neurons tuned to the expected stimulus, which in turn affects the processing of subsequent stimuli. Recent fMRI studies have revealed stimulus-specific patterns of activation in sensory cortex as a result of expectation, but this method lacks the temporal resolution necessary to distinguish pre- from post-stimulus processes. Here, we combined human MEG with multivariate decoding techniques to probe the representational content of neural signals in a time-resolved manner. We observed a representation of expected stimuli in the neural signal well before they were presented, demonstrating that expectations indeed induce a pre-activation of stimulus templates. These results suggest a mechanism for how predictive perception can be neurally implemented.

## Introduction

Perception is heavily influenced by prior knowledge^1–3^. Accordingly, many theories cast perception as a process of inference, integrating bottom-up sensory inputs and top-down expectations^4–6^. However, it is unclear how this integration is neurally implemented. It has been proposed that prior expectations lead to baseline increases in sensory neurons tuned to the expected stimulus^7–9^, which in turn leads to improved neural processing of matching stimuli^10,11^. In other words, expectations may induce stimulus templates in sensory cortex, prior to the actual presentation of the stimulus. Alternatively, top-down influences in sensory cortex may exert their influence only after the bottom-up stimulus has been initially processed, and the integration of the two sources of information may become apparent only during later stages of sensory processing^12^.

The evidence necessary to distinguish between these hypotheses has been lacking. fMRI studies have revealed stimulus-specific patterns of activation in sensory cortex as a result of expectation^9,13^, but this method lacks the temporal resolution necessary to distinguish pre- from post-stimulus periods. Here, we combined MEG with multivariate decoding techniques to probe the representational content of neural signals in a time-resolved manner^14–17^. We trained a forward model to decode the orientation of task-irrelevant gratings from the MEG signal^18,19^, and applied this decoder to trials in which participants expected a grating of a particular orientation to be presented. This analysis revealed a neural representation of the expected grating that resembled the neural signal evoked by an actually presented grating. This representation was present already before stimulus presentation, demonstrating that expectations can indeed induce the pre-activation of stimulus templates.

## Results

Participants were exposed to auditory cues that predicted the likely orientation (45° or 135°) of an upcoming grating stimulus (Fig. 1a-b). This grating was followed by a second grating that differed slightly from the first in terms of orientation and contrast. In separate runs of the MEG session, participants performed either an orientation or contrast discrimination task on the two gratings (see Methods for details).

**Figure 1.**
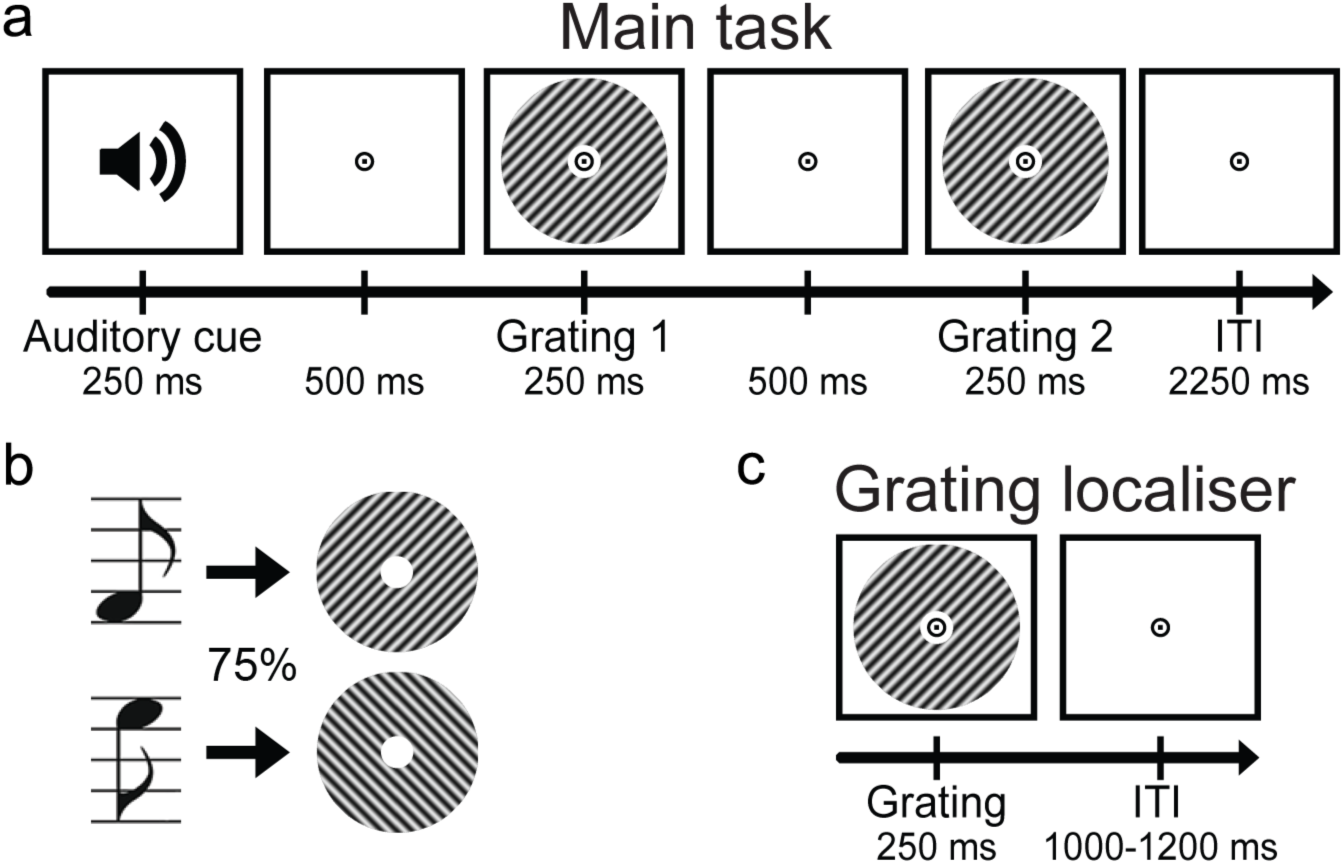
Experimental paradigm. (**a**) Each trial started with an auditory cue that predicted the orientation of the subsequent grating stimulus. This first grating was followed by a second one, which differed slightly from the first in terms of orientation and contrast. In separate runs, participants performed either an orientation or contrast discrimination task on the two gratings. (**b**) Throughout the experiment, two different tones were used as cues, each one predicting one of the two possible orientations (45° or 135°) with 75% validity. These contingencies were flipped halfway through the experiment. (**c**) In separate grating localiser runs, participants were exposed to task-irrelevant gratings while they performed a fixation dot dimming task.

### Behavioural results

Participants were able to discriminate small differences in orientation (3.9° ± 0.5°, accuracy = 74.0% ± 1.6%, mean ± sem) and contrast (4.6% ± 0.3%, accuracy = 76.6% ± 1.5%) of the cued gratings. There was no significant difference between the two tasks in terms of either accuracy (F_1,22_ = 3.38, *p* = 0.080) or reaction time (mean RT = 621 ms vs. 603 ms, *F*_1,22_ = 1.46, *p* = 0.24). Overall, accuracy and reaction times were not influenced by whether the cued grating had the expected or the unexpected orientation (accuracy: *F*_1,22_ = 0.21, *p* = 0.65; RT: *F*_1,22_ = 0.03, *p* = 0.87), nor was there an interaction between task and expectation (accuracy: *F*_1,22_ = 0.96, *p* = 0.34; RT: *F*_1,22_ = 0.42, *p* = 0.52). Note that these discrimination tasks were orthogonal to the expectation manipulation, in the sense that the expectation cue provided no information about the likely correct choice.

During the grating localiser (Fig. 1c, see Methods for details), participants correctly detected 91.2% ± 1.6% (mean ± sem) of fixation flickers, and incorrectly pressed the button on 0.2% ± 0.1% of trials, suggesting that participants were successfully engaged by the fixation task.

### MEG results – Localiser orientation decoding

As mentioned, participants were exposed to auditory cues that predicted the likely orientation of an upcoming grating stimulus. The question we wanted to answer was whether the expectations induced by these auditory cues would evoke templates of the visual stimuli prior to the presentation of the gratings. To be able to uncover such sensory templates, we trained a decoding model to reconstruct the orientation of (task-irrelevant) visual gratings (Fig. 1c) from the MEG signal, in a time-resolved manner. First, we found that this model was highly accurate at reconstructing the orientation of such gratings from the MEG signal (Fig. 2). Grating orientation could be decoded across an extended period of time (from 40 to 655 ms post-stimulus, *p* < 0.001, and from 685 to 730 ms, *p* = 0.018), peaking around 120-160 ms post-stimulus (Fig. 2c). Furthermore, in the period around 100 to 330 ms post-stimulus, orientation decoding generalised across time, meaning that a decoder trained on the evoked response at, for example, 120 ms post-stimulus could reconstruct the grating orientation represented in the evoked response around 300 ms, and vice versa (Fig. 2d). In other words, certain aspects of the representation of grating orientation were sustained over time.

**Figure 2.**
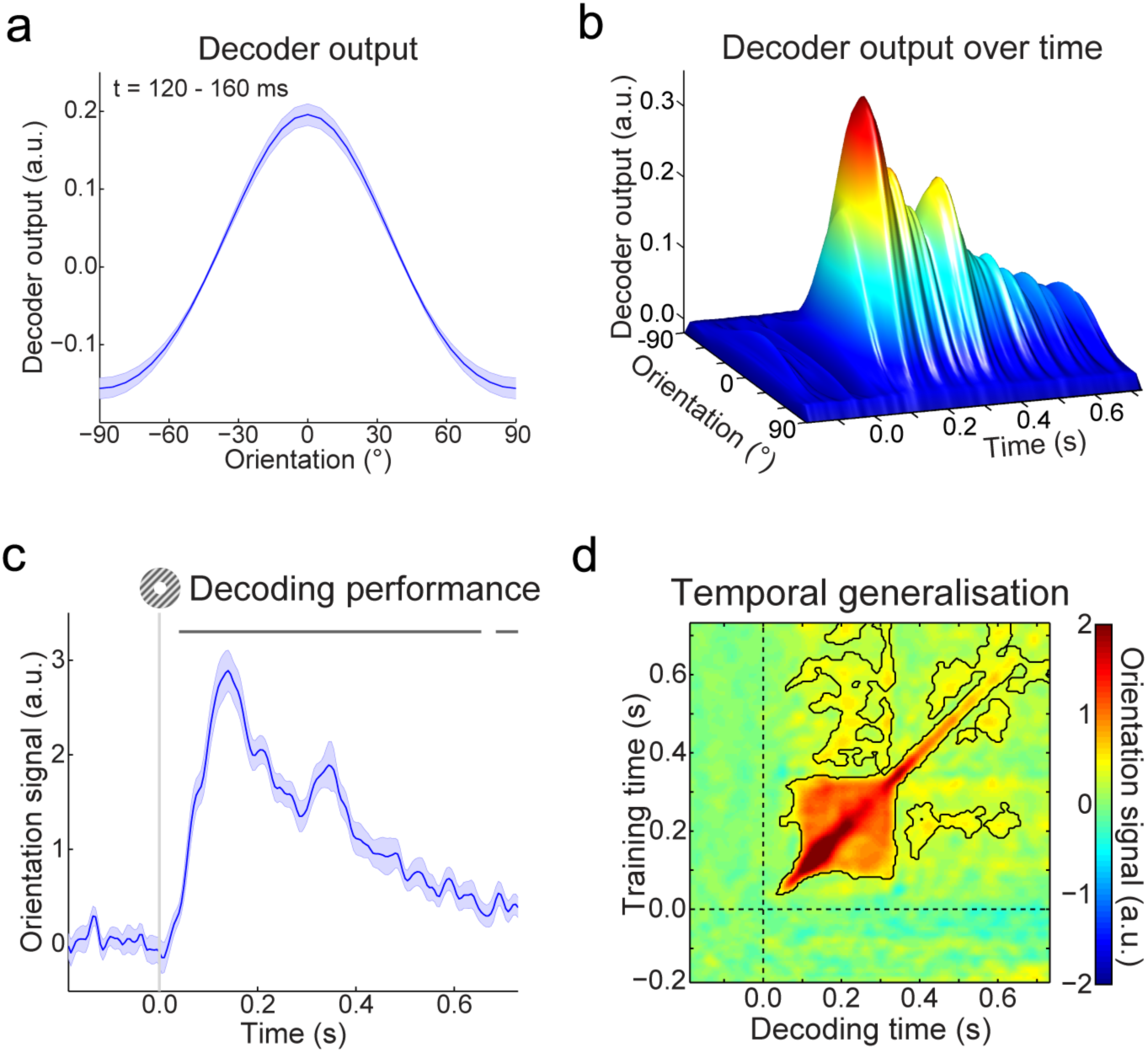
Localiser orientation decoding. (**a**) The output of the decoder consisted of the responses of 32 hypothetical orientation channels, shown here decoders trained and tested on the MEG signal 120-160 ms post-stimulus during the grating localiser (cross-validated). Shaded region represent SEM. (**b**) Decoder output over time, trained and tested in 5 ms steps (sliding window of 29.2 ms), showing the temporal evolution of the orientation signal. (**c**) The response of the 32 orientation channels collapsed into a single metric of decoding performance (see Methods), over time. Shaded region represent SEM, horizontal lines indicate significant clusters (*p* < 0.05). (**d**) Temporal generalisation matrix of orientation decoding performance, obtained by training decoders on each time point, and testing all decoders on all time points (as above, steps of 5 ms and a sliding window of 29.2 ms). This method provides insight into the sustained versus dynamical nature of orientation representations^15^. Solid black lines indicate significant clusters (p < 0.05), dashed lines indicate grating onset (t = 0s).

### MEG results – Expectation induces stimulus templates

Our main question pertained to the presence of visual grating templates induced by the auditory expectation cues during the main experiment. Therefore, we applied our model trained on task-irrelevant gratings to trials containing gratings that were either validly or invalidly predicted, respectively (Fig. 3a). In both conditions, the decoding model trained on task-irrelevant gratings succeeded in accurately reconstructing the orientation of the gratings presented in the main experiment (valid expectation: cluster from training time 60 to 410 ms and decoding time 60 to 400 ms, *p* < 0.001, and from training time 205 to 325 ms and decoding time 400 to 495 ms, *p* = 0.045; invalid expectation: cluster from training time 75 to 225 ms and decoding time 75 to 330 ms, *p* = 0.0012, and from training time 250 to 360 ms and decoding time 195 to 355 ms, *p* = 0.027).

**Figure 3.**
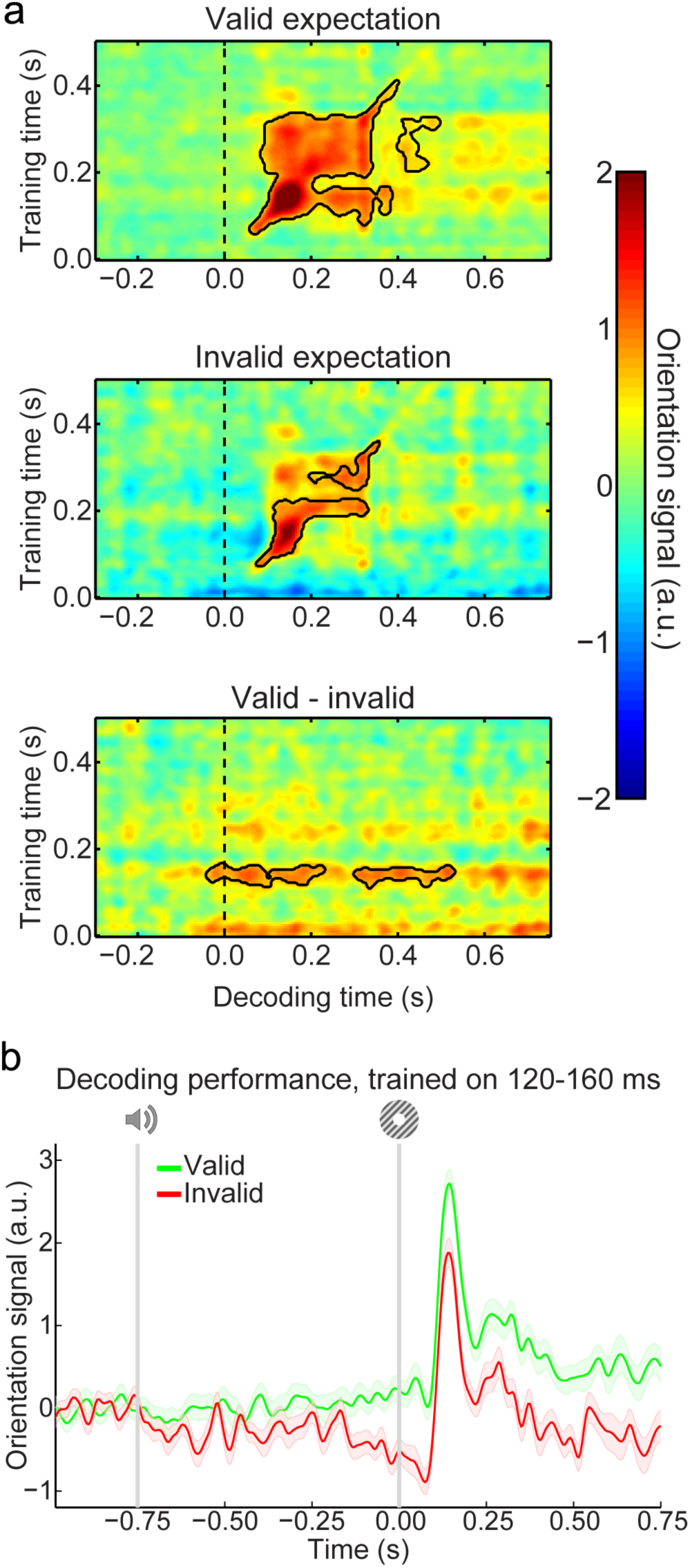
Expectation induces stimulus templates. (**a**) Temporal generalisation matrices of orientation decoding during the main experiment. Decoders were trained on the grating localiser (training time on the y-axis) and tested on the main experiment (time on the x-axis; dashed vertical line indicates t = 0s, onset of the first grating). Decoding shown separately for gratings preceded by a valid expectation (top row), invalid expectation (middle row), and the subtraction of the two conditions (i.e., the expectation cue effect, bottom row). Solid black lines indicate significant clusters (*p* < 0.05). (**b**) Orientation decoding during the main task, averaged over training time 120 – 160 ms post-stimulus during the grating localiser. That is, a horizontal slice through the temporal generalisation matrices above at the training time for which we see a significant cluster of expected orientation decoding, for visualisation. Shaded regions indicate SEM.

If the cues induced sensory templates of the expected grating, one would expect these to be revealed in the difference in decoding between valid and invalidly predicted gratings (see Material and Methods for details of the subtraction logic). Indeed, this subtraction/analyses demonstrates that the auditory expectation cues induce orientation-specific neural signals (Fig. 3a, bottom panel). These signals were present already 40 ms before grating presentation, and extended into the post-stimulus period (from decoding time -40 to 230 ms, *p* = 0.0092, and from 300 to 530 ms, *p* = 0.016). Furthermore, these signals were uncovered when the decoder was trained on around 120 to 160 ms post-stimulus during the grating localiser (Fig. 3b), suggesting that these cue-induced signals were similar to those evoked by task-irrelevant gratings. In other words, the auditory expectation cues evoked orientation-specific signals that were similar to sensory signals evoked by the corresponding actual grating stimuli.

In sum, expectations induced pre-stimulus sensory templates that influenced post-stimulus representations as well; invalidly expected gratings had to ‘overcome’ a pre-stimulus activation of the opposite orientation, while validly expected gratings were facilitated by a compatible pre-stimulus activation (Supplementary Fig. 1a). The post-stimulus carryover of these expectation signals lasted throughout the trial (Supplementary Fig. 1b).

As in previous studies using a similar paradigm^11,20^, there was no interaction between the effects of the expectation cue and the task (orientation vs. contrast discrimination) participants performed (no clusters with *p* < 0.4).

In the current study, there was no difference in the overall amplitude of the neural response evoked between validly and invalidly expected gratings (no clusters with *p* < 0.4, Supplementary Fig. 2).

## Discussion

Here, we show that expectations can induce sensory templates of the expected stimulus already before the stimulus appears. These results extend previous fMRI studies demonstrating stimulus-specific patterns of activation in sensory cortex induced by expectations, which could not resolve whether these templates indeed reflected pre-stimulus expectations, or instead stimulus specific error signals induced by the unexpected omission of a stimulus^9,13^.

The fact that expectation signals were revealed by a decoder trained on physically presented (but task-irrelevant) gratings suggests that these expectation signals resemble activity patterns induced by actual stimuli. The expectation signal remained present throughout the trial, extending into the post-stimulus period, suggesting the tonic activation of a stimulus template. These results are in line with a recent monkey electrophysiology study^10^, which showed that neurons in the face patch of IT cortex encode the prior expectation of a face appearing, both prior to and following actual stimulus presentation. When the subsequently presented stimulus is noisy or ambiguous, such a pre-stimulus template could conceivably bias perception towards the expected stimulus^21–24^.

What is the source of these cue-induced expectation signals? One candidate region is the hippocampus, which is known to be involved in encoding associations between previously unrelated, discontiguous stimuli^25^, such as the auditory tones and visual gratings used in the present study. Furthermore, fMRI studies have revealed predictive signals in the hippocampus^13,26,27^, and Reddy and colleagues^28^ reported anticipatory firing to expected stimuli in the medial temporal lobe, including the hippocampus. One intriguing possibility is that predictive signals from the hippocampus are fed back to sensory cortex^13,29,30^.

In addition to expectation, several other cognitive phenomena have been shown to induce stimulus templates in sensory cortex, such as preparatory attention^17,31^, mental imagery^32–34^, and working memory^35,36^. In fact, explicit task preparation can also induce pre-stimulus sensory templates that last into the post-stimulus period^17^. Note that in the current study the task did not require explicit use of the expectation cues, the task response was in fact orthogonal to the expectation. Furthermore, there was no difference in the expectation signal between runs in which grating orientation was task-relevant (orientation discrimination task) and when it was irrelevant (contrast discrimination task), suggestion expectation may be a relatively automatic phenomenon^11,37^. In fact, neural modulations by expectation have even been observed during states of inattention^38^, sleep^39^ and in patients experiencing disorders of consciousness^40^. One important question for future research will be to establish whether the same neural mechanism underlies the different cognitive phenomena that are capable of inducing stimulus templates in sensory cortex, or whether different top-down mechanisms are at work. Indeed, it has been suggested that expectation and attention, or task preparation, may have different underlying neural mechanisms^20,41,42^. For instance, predictive coding theories suggest that attention may modulate sensory signals in the superficial layers of sensory cortex, while predictions modulate the response in deep layers^5,43^.

One may wonder why the current study does not report a modulation of the overall neural response by expectation, while previous studies have found an increased neural response to unexpected stimuli^37,44–48^, including some using an almost identical paradigm as the current study^11,20^. Of course, the current study reports a null effect, from which it is hard to draw firm conclusions. However, it is possible that the type of measurement of neural activity plays a role in the absence of the effect. Most previous studies reporting expectation suppression in visual cortex used fMRI, while the current study used MEG. It is possible that the BOLD signal, a mass-action signal that integrates synaptic and neural activity, as well as integrating over time, is sensitive to certain neural effects that MEG, which is predominantly sensitive to synchronised activity in pyramidal neurons oriented perpendicular to the cortical surface, is not. It is even possible that within MEG, different types of sensors (i.e. magnetometers, planar and axial gradiometers) differ in their sensitivity to expectation suppression^49^.

Recent theories of sensory processing state that perception reflects the integration of bottom-up inputs and top-down expectations, but ideas diverge on whether the brain continuously generates stimulus templates in sensory cortex to pre-empt expected inputs^10,23,50,51^, or rather engages in perceptual inference only after receiving sensory inputs^52,53^. Our results are in line with the brain being proactive, constantly forming predictions about future sensory inputs. These findings bring us closer to uncovering the neural mechanisms by which we integrate prior knowledge with sensory inputs to optimise perception.

## Methods

### Participants

Twenty-three (15 female, age 26 ± 9, mean ± SD) healthy individuals participated in the experiment. All participants were right-handed and had normal or corrected-to-normal vision. The study was approved by the local ethics committee (CMO Arnhem-Nijmegen, The Netherlands) under the general ethics approval (“Imaging Human Cognition”, CMO 2014/288), and the experiment was conducted in accordance with these guidelines. All participants gave written informed consent according to the declaration of Helsinki.

### Stimuli

Grayscale luminance-defined sinusoidal grating stimuli (spatial frequency: 1.0 cycles/°) were generated using MATLAB (MathWorks, Natick, MA) in conjunction with the Psychophysics Toolbox^54^. Gratings were displayed in an annulus (outer diameter: 15° of visual angle, inner diameter: 1°), surrounding a black fixation bull’s eye (4 cd/m^2^), on a gray (580 cd/m^2^) background. The visual stimuli were presented with an LCD projector (1024 × 768 resolution, 60 Hz refresh rate) positioned outside the magnetically shielded room, and projected on a translucent screen via two front-silvered mirrors. The projector lag was measured at 36 ms, which was corrected for by shifting the time axis of the data accordingly. The auditory cue consisted of a pure tone (500 or 1000 Hz, 250 ms duration, including 10 ms on and off-ramp time), presented over MEG-compatible earphones.

### Experimental design

Each trial consisted of an auditory cue, followed by two consecutive grating stimuli (750 ms SOA between auditory and first visual stimulus) (Fig. 1a). The two grating stimuli were presented for 250 ms each, separated by a blank screen (500 ms). A central fixation bull’s eye (0.7°) was presented throughout the trial, as well as during the intertrial interval (ITI, 2250 ms). The auditory cue consisted of either a low- (500 Hz) or high-frequency (1000 Hz) tone, which predicted the orientation of the first grating stimulus (45° or 135°) with 75% validity (Fig. 1b). In the other 25% of trials, the first grating had the orthogonal orientation. Thus, the first grating had an orientation of either exactly 45° or 135°, and a luminance contrast of 80%. The second grating differed slightly from the first in terms of both orientation and contrast (see below), as well as being in antiphase to the first grating (which had a random spatial phase). The contingencies between the auditory cues and grating orientations were flipped halfway through the experiment (i.e., after four runs), and the order was counterbalanced over subjects.

In separate runs (64 trials each, ∼4.5 minutes), subjects performed either an orientation or a contrast discrimination task on the two gratings. When performing the orientation task, subjects had to judge whether the second grating was rotated clockwise or anticlockwise with respect to the first grating. In the contrast task, a judgment had to be made on whether the second grating had lower or higher contrast than the first one. These tasks were explicitly designed to avoid a direct relationship between the perceptual expectation and the task response. Subjects indicated their response (response deadline: 750 ms after offset of the second grating) using an MEG-compatible button box. The orientation and contrast differences between the two gratings were determined by an adaptive staircase procedure^55^, being updated after each trial. This was done to yield comparable task difficulty and performance (∼ 75% correct) for the different tasks. Staircase thresholds obtained during one task were used to set the stimulus differences during the other task, in order to make the stimuli as similar as possible in both contexts. As in previous studies using a similar paradigm^11,20^, there was no interaction between the effects of the expectation cue and the task participants performed, and therefore we collapsed over the two tasks in all MEG analyses.

All subjects completed eight runs (four of each task, alternating every two runs, order was counterbalanced over subjects) of the experiment, yielding a total of 512 trials. The staircases were kept running throughout the experiment. Before the first run, as well as in between runs four and five, when the contingencies between cue and stimuli were flipped, subjects performed a short practice run containing 32 trials of both tasks (∼4.5 minutes).

Interleaved with the main task runs, subjects performed eight runs of a grating localiser task (Fig. 1c). Each run (∼2 min) consisted of 80 grating presentations (ITI uniformly jittered between 1000 and 1200 ms). The grating annuli were identical to those presented during the main task (80% contrast, 250 ms duration, 1.0 cycles/°, random spatial phase). Each grating had one of eight orientations (spanning the 180° space, starting at 0°, in steps of 22.5°), each of which was presented ten times per run in pseudorandom order. A black fixation bull’s eye (4 cd/m^2^, 0.7° diameter, identical to the one presented during the main task runs) was presented throughout the run. On 10% of trials (counterbalanced across orientations), the black fixation point in the centre of the bull’s eye (0.2°, 4 cd/m^2^) briefly turned gray (324 cd/m^2^) during the first 50 ms of grating presentation. Participants task was to press a button (response deadline: 500 ms) when they perceived this fixation flicker. This simple task was meant to ensure central fixation, while rendering the gratings task-irrelevant. Trials containing fixation flickers were excluded from further analyses.

Finally, participants were exposed to a tone localiser (∼1.5 min), presented at the start, end, and halfway through the MEG session. These runs consisted of 81 presentations of the two tones used in the main experiment. Data from these runs were not analysed further.

Prior to the MEG session (1–3 days), all participants completed a behavioural session. The aim of this session was to familiarise participants with the tasks and to initialise the staircase values for both the orientation and the contrast discrimination task (see above). The behavioural session consisted of written instructions and 32 practice trials of each task, followed by four runs (∼4.5 min each) of the main experiment (each task twice, alternating between runs, cue contingencies switching between the second and third run). Finally, participants were exposed to one run each of the grating and tone localiser, to familiarise them with the procedure.

### MEG recording and preprocessing

Whole-head neural recordings were obtained using a 275-channel MEG system with axial gradiometers (CTF Systems, Coquitlam, BC, Canada) located in a magnetically shielded room. Throughout the experiment, head position was monitored online, and corrected if necessary, using three fiducial coils that were placed on the nasion and on earplugs in both ears ^56^. If subjects had moved their head more than 5 mm from the starting position they were repositioned during block breaks. Furthermore, both horizontal and vertical electrooculograms (EOGs), as well as an electrocardiogram (ECG) were recorded to facilitate removal of eye- and heart-related artifacts. The ground electrode was placed at the left mastoid. All signals were sampled at a rate of 1200 Hz.

The data were preprocessed offline using FieldTrip^57^ (www.fieldtriptoolbox.org). In order to identify artifacts, the variance (collapsed over channels and time) was calculated for each trial. Trials with large variances were subsequently selected for manual inspection and removed if they contained excessive and irregular artifacts. Independent component analysis was subsequently used to remove regular artifacts, such as heartbeats and eye blinks. Specifically, for each subject, the independent components were correlated to both EOGs and the ECG to identify potentially contaminating components, and these were subsequently inspected manually before removal. For the main analyses, data were low-pass filtered using a two-pass Butterworth filter with a filter order of 6 and a frequency cutoff of 40 Hz. To rule out that the temporal smoothing caused by low-pass filtering may have artificially decreased the onset latency of neural signals, we repeated the decoding analyses (see below) on data that were not low-pass filtered (Supplementary Fig. 3). Here, only notch filters were applied at 50, 100 and 150 Hz to remove line noise and its harmonics. Finally, main task data were baseline corrected on the interval of –250 to 0 ms relative to auditory cue onset, and grating localiser data were baseline corrected on the interval of -200 to 0 ms relative to visual grating onset.

### Event-related field analysis

Event-related fields (ERFs) were calculated per participant, and subjected to a planar gradient transformation^58^ before averaging across participants. The planar transformation simplifies the interpretation of the sensor-level data because it typically places the maximal signal above the source. To avoid differences in the amount of noise when comparing conditions with different numbers of trials, we matched the trial count by randomly selecting a subsample of trials from the conditions with more trials (i.e., valid expectations).

### Orientation decoding analysis

To probe sensory representations in the visual cortex, we used a forward modelling approach to reconstruct the orientation of the grating stimuli from the MEG signal^17–19,59^. The forward modelling approach was two-fold. First, a theoretical forward model was postulated that described the measured activity in the MEG sensors, given the orientation of the presented grating. Second, this forward model was used to obtain an inverse model that specified the transformation from MEG sensor space to orientation space. The forward and inverse models were estimated on the basis of the grating localiser data. The inverse model was then applied to the data from the main experiment, in order to generalise from sensory signals evoked by task-irrelevant gratings to the gratings and expectation signals evoked in the main task. To test the performance of the model we also applied it to the localiser data itself, using a cross-validation approach in which in each iteration one trial of each orientation was used at the test set, and the remaining data were used as the training set.

The forward model was based on work by Brouwer and Heeger^18,19^ and involved 32 hypothetical channels, each with an idealised orientation tuning curve. Each channel consisted of a half-wave-rectified sinusoid raised to the fifth power, and the 32 channels were spaced evenly within the 180° orientation space, such that a tuning curve with any possible orientation preference could be expressed exactly as a weighted sum of the channels. Arranging the hypothesised channel activities for each trial along the columns of a matrix **C** (32 channels × *n* trials), the observed data could be described by the following linear model: 
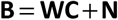
where **B** are the (*m* sensors × *n* trials) MEG data, **W** is a weight matrix (*m* sensors × 32 channels) that specifies how channel activity is transformed into sensory activity, and **N** are the residuals (i.e., noise).

In order to obtain the inverse model, we estimated an array of spatial filters that, when applied to the data, aimed to reconstruct the underlying channel activities as accurately as possible. In doing so, we extended Brouwer and Heeger’s^18,19^ approach in three respects. First, since the MEG signal in (nearby) sensors is correlated, we took into account the correlational structure of the noise. Second, we estimated a spatial filter for each orientation channel independently. As a result, the number of channels used in our model was not constrained, whereas the maximum number of channels would otherwise be dependent on the number of presented orientations. In practice, this resulted in smoothing in orientation space, because the channels were not truly independent. Third, each filter was normalised such that the magnitude of its output matched the magnitude of the underlying channel activity it was designed to recover. Prior to estimating the inverse model, **B** and **C** were demeaned such that their average over trials equalled zero, for each sensor and channel, respectively.

As stated above, the inverse model was estimated on the basis of the grating localiser data. On each localiser trial, one of eight orientations was presented (see above), and the hypothetical responses of each of the channels could thus be calculated for each trial, resulting in the response row vector **c***_train,i_*, of length *n_train_* trials, for each channel *i*. The weights on the sensors **w**_*i*_ could now be obtained through least squares estimation, for each channel:

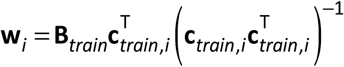
where **B***_train_* are the (*m* sensors × *n_train_* trials) localiser MEG data. Subsequently, the optimal spatial filter **v**_*i*_ to recover the activity of the *i*-th channel was obtained as follows^16^: 
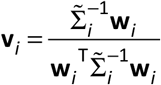
Where 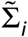 is the regularised covariance matrix for channel *i*. Incorporating the noise covariance in the filter estimation leads to the suppression of noise that arises from correlations between sensors. The noise covariance was estimated as follows: 
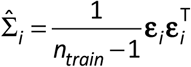

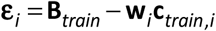
where *n_train_* is the number of training trials. For optimal noise suppression, we improved this estimation by means of regularization by shrinkage, using the analytically determined optimal shrinkage parameter (for details, see^60^), yielding the regularised covariance matrix 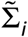.

Such a spatial filter was estimated for each hypothetical channel, yielding an *m* sensors × 32 channel filter matrix **V**. Given that we performed our decoding analysis in a time-resolved manner, **V** was estimated at each time point of the training data, in steps of 5 ms, resulting in array of filter matrices, or decoders. To improve the signal-to-noise ratio, the data were first averaged within a window of 29.2 ms centred on the time point of interest. The window length of 29.2 ms was based on an a priori chosen length of 30 ms, but minus one sample such that the window contained an odd number of samples for symmetric centring^16^. These filter matrices could now be applied to estimate the orientation channel responses in independent data – in this case, the trials from the main experiment: 
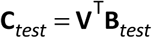
where **B***_test_* are the (*m* sensors × *n_test_* trials) main experiment data. These channel responses were estimated at each time point of the test data, in steps of 5 ms, with the data being averaged within a window of 29.2 ms at each step. This procedure resulted in a four-dimensional (training time × testing time × 32 channel × *n_test_*) matrix of estimated channel responses for each trial in the main experiment.

Each trials’ channel responses were shifted such that the channel with its hypothetical peak response at the orientation presented on that trial (i.e. 45° or 135°) ended up in the position of the 0° channel, before averaging over trials within each condition (i.e., valid vs. invalid expectation). Thus, the presented orientation was defined as 0°, by convention. Note that for 3D surface plots that show the evolution of channel responses over time (e.g., Fig. 2b), the response of the 90° channel (i.e., orthogonal to the presented orientation) was used as a baseline, to avoid negative numbers for visualisation purposes.

To quantify decoding performance, the channel responses for a given condition were converted into polar form and projected onto a vector with angle 0° (the presented orientation, see above).

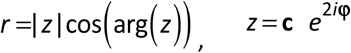
where **c** is a vector of estimated channel responses, and **φ** is the vector of angles at which the channels peak (multiplied by 2 to project the 180° orientation space onto the full 360° space). The scalar projection *r* indicates the strength of the decoder signal for the orientation presented on screen. (Note that subtracting the estimated response of the 90° channel from that of the 0° channel yielded virtually identical results, data not shown.) This quantification yielded (training time × testing time) temporal generalisation matrices of orientation decoding performance.

In order to isolate any orientation-specific neural signals evoked by the expectation cues, we applied the following subtraction logic. On valid expectation trials, the expected and presented orientations are identical, and thus the orientation signal induced by both the cue and stimulus be expected to be positive, by convention. On invalid expectation trials on the other hand, the expected and presented orientations are orthogonal, and thus the orientation signal induced by the stimulus would be positive and the signal induced by cue would be expected to be negative. Thus, subtracting the orientation decoding signal on invalid trials from that on valid trials would subtract out the stimulus-evoked signal while revealing any cue-induced orientation signal.

### Statistical testing

Neural signals evoked by the different conditions were statistically tested using nonparametric cluster-based permutation tests^61^. For ERF analyses, we averaged over the spatial (sensor) dimension, on the basis of independent localisation of the 10 sensors that showed the strongest visual-evoked activity during the grating localiser between 50 and 150 ms post-stimulus. Therefore, our statistical analysis considered one-dimensional (temporal) clusters. For orientation decoding analyses, the data consisted of two-dimensional (training time × testing time) decoding performance matrices, and the statistical analysis thus considered two-dimensional clusters. For both one-and two-dimensional data, univariate t-statistics were calculated for the entire matrix and neighbouring elements that passed a threshold value corresponding to a *p*-value of 0.01 (two-tailed) were collected into separate negative and positive clusters. Elements were considered neighbours if they were directly adjacent, either cardinally or diagonally. Cluster-level test statistics consisted of the sum of *t*-values within each cluster, and these were compared to a null distribution of test statistics created by drawing 10,000 random permutations of the observed data. A cluster was considered significant when its *p*-value was below 0.05 (two-tailed).

## Acknowledgements

This work was supported by The Netherlands Organisation for Scientific Research (NWO) to P.K. (Rubicon grant 446-15-004) and F.d.L. (VIDI grant 452-13-016) and the James S McDonnell Foundation to F.d.L. (Understanding Human Cognition 220020373). The authors would like to thank Mariya Manahova for data collection.

## Author contributions

P.K. and F.d.L. designed the experiment, P.K. and P.M. conducted the experiment, P.K. and P.M. analysed the data, P.K., P.M. and F.d.L. wrote the paper.

